# Developmental changes of opsin gene expression in ray-finned fishes (Actinopterygii)

**DOI:** 10.1101/2022.04.28.489877

**Authors:** Nik Lupše, Monika Kłodawska, Veronika Truhlářová, Prokop Košátko, Vojtěch Kašpar, Arnold Roger Bitja Nyom, Zuzana Musilova

**Affiliations:** Department of Zoology, Faculty of Science, Charles University, Vinicna 7, 12844 Prague, Czech Republic; University of South Bohemia in České Budějovice, Faculty of Fisheries and Protection of Waters, South Bohemian Research Center of Aquaculture and Biodiversity of Hydrocenoses, Research Institute of Fish Culture and Hydrobiology, Zátiší 728/II, 389 25 Vodňany, Czech Republic; Department of Management of Fisheries and Aquatic Ecosystems, University of Douala, Douala, Cameroon; Department of Biological Sciences, University of Ngaoundéré, Ngaoundéré, Cameroon

**Keywords:** opsin, evolution, Actinopterygii, development, vision, gene expression

## Abstract

Fish often change their habitat and trophic preferences during development. Dramatic functional differences between embryos, larvae, juveniles and adults also concern sensory systems, including vision. Here we focus on the photoreceptors (rod and cone cells) in the retina and their gene expression profiles during the development. Using comparative transcriptomics on 63 species, belonging to 23 actinopterygian orders, we report general developmental patterns of opsin expression, mostly suggesting an increased importance of the rod opsin (*RH1*) gene and the long-wavelength sensitive (*LWS*) cone opsin, and a decreasing importance of the shorter wavelength sensitive cone opsin throughout development. Furthermore, we investigate in detail ontogenetic changes in 14 selected species (from Polypteriformes, Acipenseriformes, Cypriniformes, Aulopiformes and Cichliformes), and we report examples of expanded cone opsin repertoires, cone opsin switches (mostly within *RH2*) and increasing rod:cone ratio as evidenced by the opsin and phototransduction cascade genes. Our findings provide molecular support for developmental stage-specific visual palettes of ray-finned fishes and shifts between, which most likely arose in response to ecological, behavioural and physiological factors.

## INTRODUCTION

Fish visual systems are very diverse, and they vary in morphology, physiology and spectral sensitivity (Hunt et al. 2014, Carleton et al. 2020, Musilova et al. 2021). Vertebrate vision is enabled by cone and rod photoreceptors in the retina, which carry light-sensitive molecules composed of an opsin protein bound to a light absorbing, vitamin A-derived chromophore (Lamb 2013). In fishes, there are usually four types of cone opsins [*SWS1* and *SWS2;* commonly found in single cones, whereas *RH2* and *LWS* in double cones; with the respective peak sensitivity ranges of 347 – 383 nm, 397 – 482 nm, 452 – 537 nm and 501 – 573 nm; (Carleton et al. 2020)] used for photopic and colour vision, and one rod opsin (rhodopsin, *RH1* or Rho) for scotopic vision in dim-light conditions (Carleton et al. 2020). Through gene duplications followed by functional diversifications, extant teleost fishes reached a median of seven cone opsin genes within their genomes (Musilova et al. 2019a). Throughout the phylogeny, teleost genomes contain more copies of double-cone genes (middle and longer-wavelength sensitive; *RH2* and *LWS*) than that of single-cones (shorter-wavelength *SWS1* and *SWS2*). While the *SWS1* is often missing from the genome or seen in one, at best two copies (Musilova et al, 2021) and *SWS2* seen in up to three copies (Cortesi et al., 2015), teleost genomes can contain up to eight copies of *RH2* (Musilova & Cortesi, 2021) and up to five copies of *LWS* (Cortesi et al., 2021). Unlike cone opsins, rod opsin duplicates are rarely found, most often in mesopelagic lineages (Pointer et al. 2007, Musilova et al. 2019a, Lupše et al. 2021). Higher copy number is considered beneficial by providing more “substrate” for selection, as well as for alternative gene expression of the variants within the opsin type.

The formation of the eye, and expression of opsin genes, starts already at the embryonic stage (Hagedorn and Fernald 1992, Carleton et al. 2008). Still, eyes continue to grow, and new photoreceptors are being added throughout life (Fernald 1985). Within the retina, cone photoreceptors are first to develop, followed by temporally and spatially distinct rods (Raymond 1995, Helvik et al. 2001, Shen & Raymond 2004). For example, in zebrafish, photoreceptor progenitor cells start out by first differentiating into cones before rods are added later during development (Sernagor et al. 2006), suggesting that vision changes with age. This cone-to-rod developmental sequence is likely shared across vertebrates (Atlantic cod: Valen et al. 2016; zebrafish: Sernagor et al. 2006; mice: Mears et al. 2001; rhesus monkey: La Vail et al. 1991) and appears to hold even for teleost species with an all-rod retina in the adult stage (Lupše et al. 2021).

Photic conditions can change spatially and temporally, resulting in a visually heterogeneous environment in which visual systems of fishes are expected to be under natural selection that favours those that match the local environment (Carleton et al. 2016). For example, longer and shorter wavelengths are scattered and filtered out with increasing water depth and consequently, fishes living in deep-water habitats such as sculpins of Lake Baikal (Hunt 1997), cichlids of lakes Malawi and Tanganyika (Sugawara et al. 2005; Ricci et al. 2022), and African crater lakes (Malinsky et al. 2015; Musilova et al. 2019b), as well as deep-sea fishes (Douglas et al. 2003, Lupše et al. 2021) have visual systems sensitive to the blue-green part of the visible spectrum. Adaptation can be achieved either through functional diversifications of opsin genes when mutations at key-spectral tuning sites shift the peak spectral sensitivity (λ_max_) of the photopigment (Yokoyama 2008, 2020), or by regulation of the opsin gene expression. This can be achieved when a subset of opsin genes is expressed and altered among or within species and even within the same individuals during ontogeny (Carleton & Kocher 2001, Manousaki et al. 2013, Carleton 2016, Lupše et al. 2021).

Before reaching the juvenile or sexually mature adult stage, fish larvae undergo major anatomical, physiological, behavioural and quite often, ecological changes (Evans & Browman 2004, Carleton et al. 2020). Developmental shift in habitat preference is often suggested to drive ontogenetic changes in opsin expression (e.g. cichlids: Carleton et al. 2016, Härer et al. 2017; black bream: Shand et al., 2002; eel: Cottrill et al. 2009; squirrelfishes and soldierfishes: Fogg et al. 2021; clown anemonefish: Roux et al. 2022; damselfishes: Stieb et al. 2016, bluefin killifish: Chang et al. 2021; gambusia: Chang et al. 2020; rainbow trout: Allison et al. 2006; dottybacks: Cortesi et al. 2016; starry flounder (Savelli et al. 2018); deep-sea fishes: de Busserolles et al. 2014, Lupše et al. 2021). However, habitat-related changes of photic conditions solely do not always result in different and stage-specific visual system modifications, as seen in the Atlantic cod (Valen et al. 2016) or the spotted unicornfish (Tettamanti et al. 2019). Shifts in diet (planktivory, carnivory, herbivory) and activity patterns (diurnal, nocturnal, crepuscular) (King and McFarlane 2003; Helfman et al. 2009, Fogg et al. 2021), in addition to developmental or phylogenetic constraints seem to also play a role in shaping the visual diversity of fishes and potential age-related shifts of it.

Here we aim to investigate ontogenetic changes of the opsin and phototransduction cascade gene expression across ray-finned fishes, to estimate presence and relative abundance of opsin gene classes, and to elucidate general and/or taxon-specific patterns. For the purpose of this study we have sequenced and analysed 1) retinal transcriptomes of different developmental stages of 14 species, belonging to five major actinopterygian orders: *Polypterus senegalensis* (Polypteriformes), *Acipenser ruthenus* (Acipenseriformes), *Abramis brama* and *Vimba vimba* (both Cypriniformes), *Scopelarchus* spp. and *Coccorella atlantica* (both Aulopiformes), *Coptodon bemini*, *C. imbriferna*, *C. flava*, *C. snyderae*, *C. thysi*, *Sarotherodon linnellii*, *S. lohbergeri* and *Stomatepia pindu* (all Cichliformes from the Bermin and Barombi Mbo lakes). 2) We have complemented this data set by publicly available embryonic/larval/juvenile/adult transcriptomes belonging to 49 species and 21 orders, some of which have never been analysed for visual gene expression before. In total, the comprehensive data set of 63 species from 23 ray-finned fish orders allows us to focus on development of the opsin gene expression, and rod and cone cell identity throughout actinopterygian evolution.

## METHODS AND MATERIALS

### Data and sample collection

Transcriptomes belonging to taxa deemed as focal groups, which were inspected for age-specific copies and presented in detail in Figure 3, were obtained from specimens (N=73) caught solely for the purpose of this study. In detail, 16 specimens were classified as larvae, 4 as juveniles, 3 as subadults and 50 as adults (Figure 3, Supp Table). *Polypterus senegalensis* larvae were collected in the rearing facility of the Department of Zoology, Charles University, and the adults were purchased from the aquarium trade. *Acipenser ruthenus* and *Abramis brama* were collected at the rearing facility in Vodňany, and in local water bodies (adults: Velky Tisy pond, Klicava dam, Lipno dam; larvae: Vltava and Elbe rivers), Czech Republic, respectively. Both mesopelagic taxa, *Scopelarchus* spp. and *Coccorella atlantica*, were collected in the Sargasso Sea and originate from Lupše et al. (2021). Crater lake cichlids were collected in lakes Barombi Mbo and Bermin (Cameroon, West Africa) between 2013 and 2018 (research permit numbers: 0000047,49/MINRESI/B00/C00/C10/nye, 0000116,117/MINRESI/ B00/C00/C10/C14, 000002-3/MINRESI/B00/C00/C10/C11, 0000032,48-50/MINRESI/B00/C00/C10/C12). Larvae were caught by fine-meshed nets and fixed in RNAlaterTM immediately. Adults were collected using gill nets and selective capturing by snorkelling in the shallow-water zone. For all species, fin clips were taken from specimens and stored in 96% EtOH for sub-sequent molecular analyses. Larval samples were fixed in RNAlaterTM (ThermoFisher) and stored at −80 °C until further use. Adults of all species were euthanised on site with eyes or retinae extracted, fixed in RNAlaterTM and stored at −80 °C upon arrival to the laboratory.

To obtain publicly available transcriptomes used in this study (Fig 1, Supp Table), we searched the largest publicly available repository of high throughput sequencing data, the Sequence Read Archive (SRA), using the following topic search term: ‘*(embryo* OR larva* OR juvenile* OR adult*) AND (retina* OR eye* OR head* OR whole*) AND (taxon name * OR fish*)*’. Whenever possible, we have analysed up to three specimens per stage per species (Fig 1, Supp Table). In case of embryos, specimens closest to hatching were analysed. The entire dataset analysed, including de-novo transcriptomes described below, includes 215 samples of which, based on morphology, 56 were classified as embryos, 40 as larvae, 25 as juveniles, 3 as subadults and 91 as adults (Figs1 and 3, Supp Table). Sample IDs, number of raw reads, individual accession numbers for BioProject PRJNA841439 and further parameters are listed in the Supplementary Table.

**Fig. 1:**
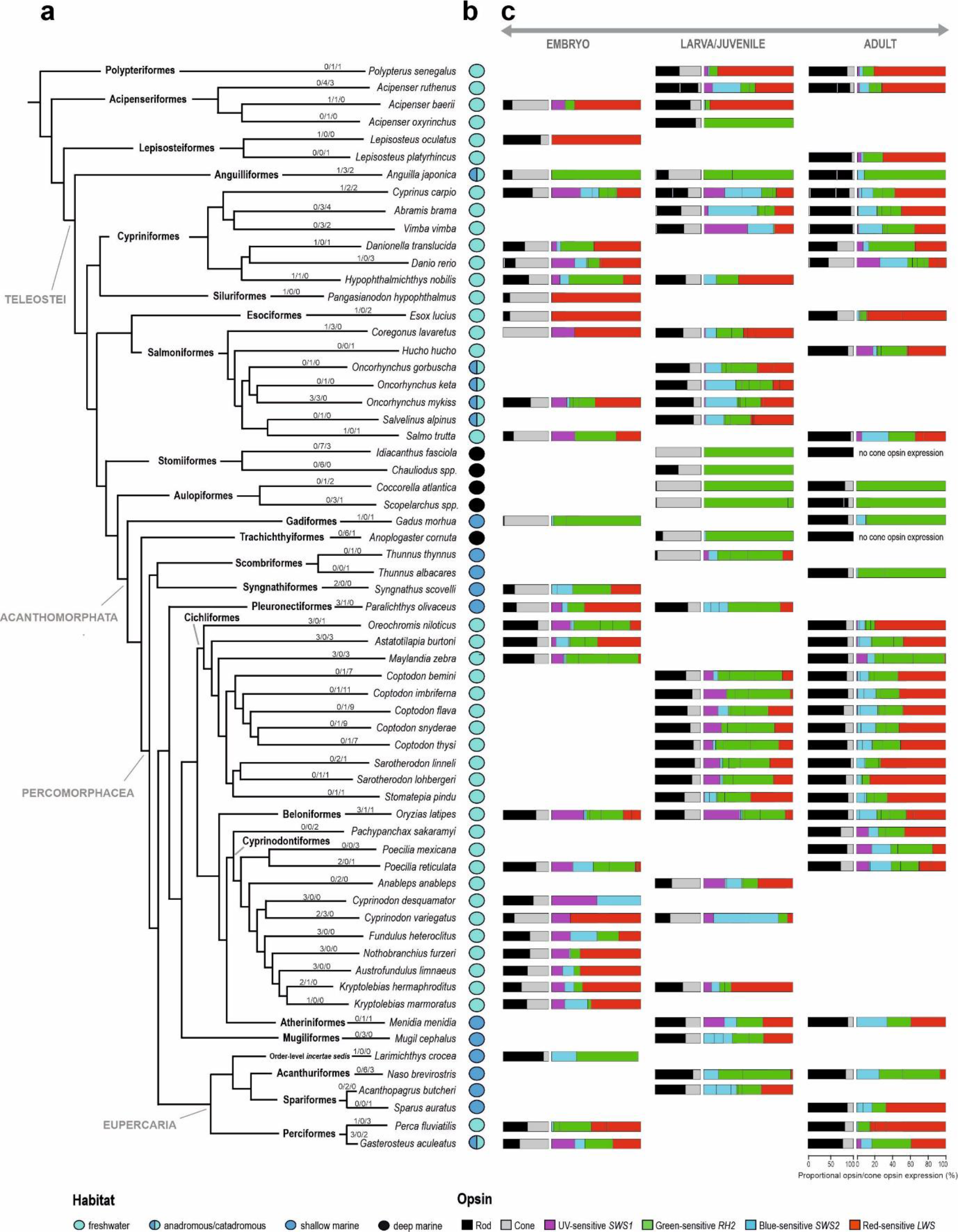
Opsin gene expression in different developmental stages of ray-finned fishes (Actinopterygii). (A) Simplified phylogeny of the 63 species, belonging to 23 orders, for which the transcriptomes were analysed (topology after Betancur et al. 2017). Numbers above branches represent number of individuals per developmental stage analysed (embryo/larva+juvenile/adult). (B) Information on habitat preference, obtained from https://www.fishbase.de. Separation between the shallow and deep marine species is 200m. Information on depth obtained from https://obis.org/. (C) Proportional opsin gene expression (horizontal bars) at different developmental stages. First (shorter) bar represents mean proportional expression of rod and cone opsins. Cone opsin expression (grey) is depicted as the sum of the expression of all four classes of cone opsin genes (SWS1, SWS2, RH2, and LWS). If several rod opsin genes (black) were expressed, the different proportions of their expression are distinguished with white vertical bars. Second (longer) bar represents mean proportional expression of different cone opsins. Black vertical bars within gene classes separate different copies, if co-expressed. For details, see Supplementary Table.

### Transcriptome sequencing and analyses

Total RNA was extracted from the whole eyes or retinal tissue using either the RNeasy micro or mini kit (Qiagen). The extracted RNA concentration and integrity were verified on a 2100 Bioanalyzer (Agilent) and Qubit Fluorometer (Thermofisher Scientific). RNAseq libraries were constructed in-house from unfragmented total RNA using Illumina’s NEBNext Ultra II Directional RNA library preparation kit, NEBNext Multiplex Oligos and the NEBNext Poly(A) mRNA Magnetic Isolation Module (New England Biolabs). Multiplexed libraries were sequenced on the Illumina HiSeq 2500 platform as 150 bp paired-end (PE) reads. The sequence data was quality-checked using FastQC (Andrews 2017). Opsin gene expression was then quantified using Geneious software version 11.0.3 (Kearse et al. 2012). In case of each sample, we mapped the reads against a general genomic reference dataset for all visual opsin genes composed of Nile tilapia, zebrafish and the longnose gar, using the Medium-sensitivity settings in Geneious. This enabled us to capture most of the cone and rod opsin specific reads and create species-specific opsin references. If needed, paralogous genes were subsequently disentangled following the methods in Musilova et al. (2019a) and de Busserolles et al. (2017). Transcriptome reads were then re-mapped to the newly created (species-specific) references with Medium-Low sensitivity to obtain copy-specific expression levels. We report opsin gene proportional expression in relation to the total opsin gene expression which was calculated using FPKM (Fragments Per Kilobase of transcript Per Million reads), taking into account the library size, the length of each gene and number of mapped reads (Supp Table). The abovementioned quantification of opsin gene expression was also used on transcriptomes obtained from SRA. Identical pipeline was used for quantification of *GNAT1*/*2* genes in selected taxa (Fig 3).

### Statistical analyses

To formally test whether opsin gene expression differs between developmental stages, we applied the beta regression models specifically designed to analyse the proportional data sets and percentages. We used the R package betareg (Cribari-Neto & Zeileis 2010), which allows handling of non-transformed data. The beta distribution has a highly flexible shape and is, hence, suitable to fit the dependent variable (in our case the proportional expression of each opsin gene) in the standard unit interval (0,1) with a mean related to a set of categorical regressors (in our case developmental stage). We tested the difference for each cone opsin gene class separately (i.e, *SWS1*, *SWS2*, *RH2* and *LWS*), then for the sum of single cone-(*SWS1*+*SWS2*), and double cone opsins (*RH2*+*LWS*), and additionally also for rods (*RH1*) and cones (*SWS1*+*SWS2*+*RH2*+*LWS*).

## RESULTS AND DISCUSSION

### General developmental patterns of opsin gene expression across the actinopterygian phylogeny – cone-to-rod developmental constraint

The analysis of the opsin gene expression in 63 ray-finned fishes revealed that, generally, the ratio of the rod opsin (*RH1* or Rho, λ_max_: 447–525 nm) to cone opsin expression increases with age in analysed species (Figs 1 and 2, Table 1, Supp Table; p = <0.001). This is in accord with the cone-to-rod development of the retina which starts with cone cells, and rods appearing only later (Sernagor et al. 2006, Valen et al. 2016, Lupše et al. 2021). The increasing rod:cone cell ratio is further confirmed by the expression of the phototransduction cascade gene *GNAT1* (rod specific) vs. *GNAT2* (cone specific), Fig. 3b. Rod opsin and *GNAT1*/*2* usage increases significantly already during the larval and juvenile stage, before finally transforming into sexually mature adults with rod-dominant retina (Figs 1 and 2, Supp Table). It thus seems that larval vision is mostly driven by cone vision, while the ability to perform well in low-light conditions appears consequently, at later developmental stages (Evans and Browman 2004, Evans and Fernald 1990). Functionally, rods generally allow for an improvement in visual acuity and startle responses in fishes (Fuiman 1993, Pankhurst et al. 1993, Fuiman and Delbos 1998) and are also associated with motion sensitivity and the appearance of novel behaviours, such as schooling (Hunter & Coyne 1982). More specifically, higher rod expression increases individual performance of fishes living in the deep-sea (de Busserolles et al. 2020, Lupše et al. 2021). Additionally, laboratory experiments have shown that the ability to follow a rotating stripe pattern (the optomotor drum) might be dependent on rod formation and retinal development, as it is not seen in stages or specimens lacking rods (Blaxter 1986, Carvalho et al. 2002, Magnuson et al. 2020).

**Fig. 2:**
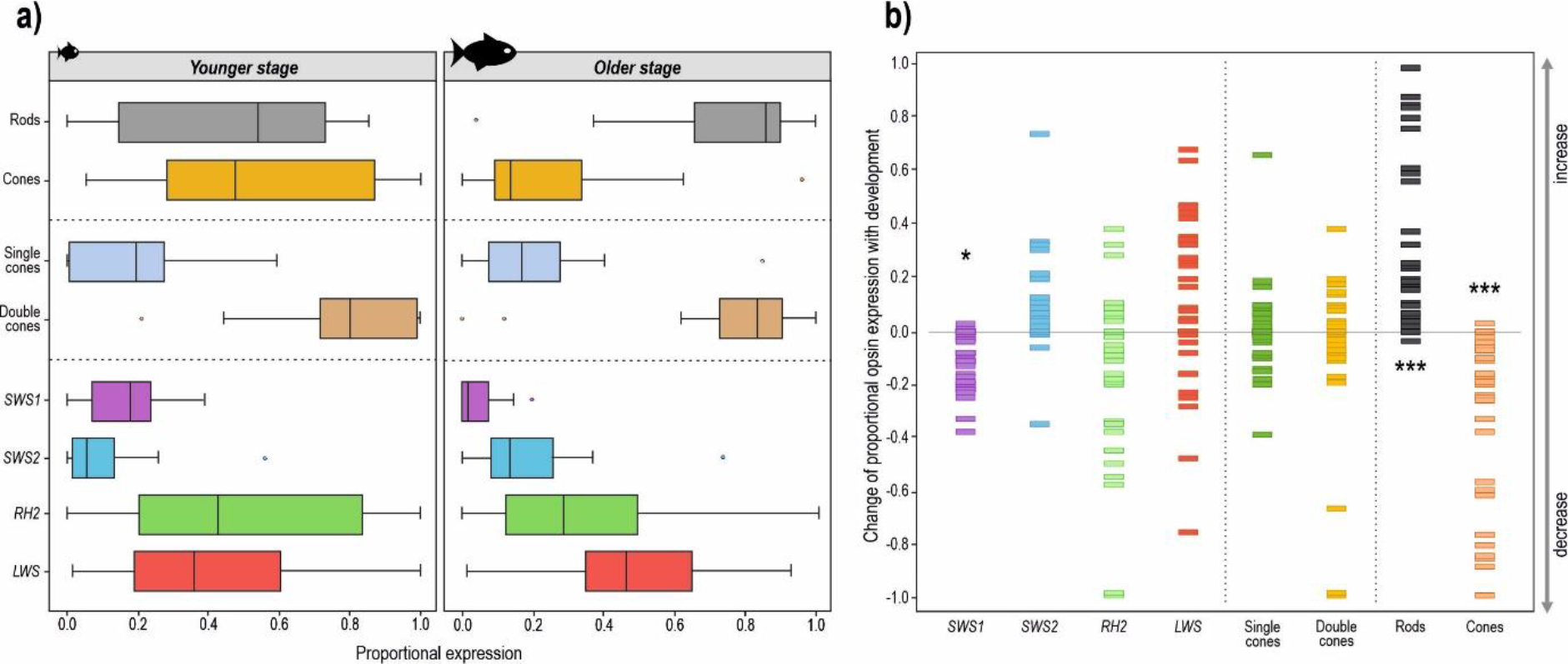
General patterns of age-related opsin expression changes. (A) Interquartile ranges (25^th^ and 75^th^ percentiles) and whiskers show data dispersion (proportional expression) across different opsins for the youngest and oldest analysed stage. Data medians are presented as solid vertical lines. To avoid over-representation of certain taxa (e.g. five Coptodon species), data points (N=32) represent mean genus values, and are comprised only of species that had at least two developmental stages analysed. (B) Change of opsin expression (positive/negative) with development, calculated as a difference between the mean opsin expression in the oldest and the youngest stage of a certain genus. Resulting values are represented by rectangles (N=32), centered at the mean. Lower half of the plot (values below 0.0) shows a decrease, and the upper half (values above 0.0) an increase in proportional expression with age. Significant differences found by beta regression models are marked by asterisks (see Table 1 for details).

**Fig. 3:**
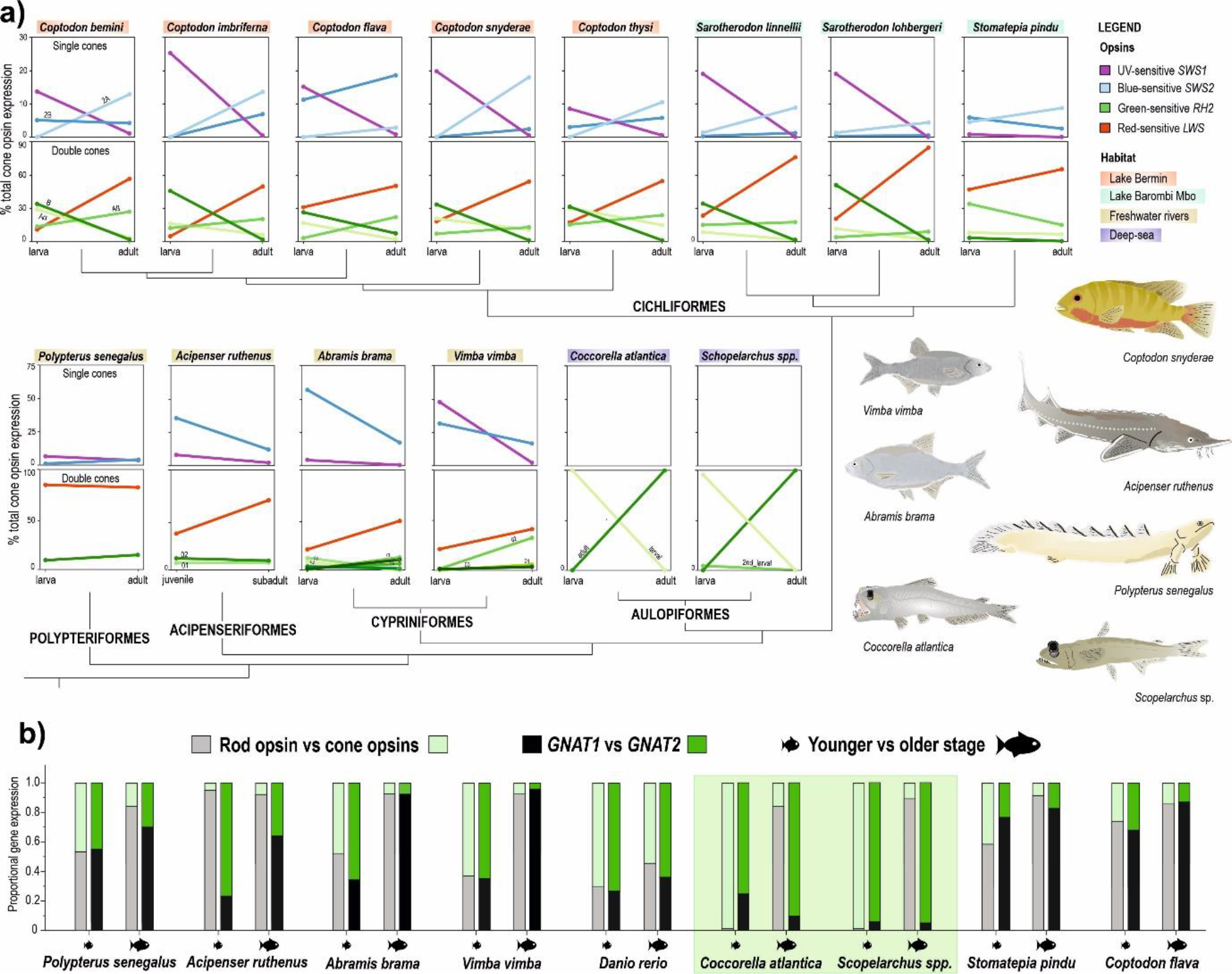
Cone opsin gene switches, age-specific copies and phototransduction cascade gene expression of representative taxa specifically sequenced for this study. (A) Detailed presentation of ontogenetic changes of opsin expression in selected polypteriform, acipenseriform, cypriniform, aulopiform and cichliform species. Interconnected dots are coloured according to specific single and double cone opsins and present mean expression values for specific developmental stages. In cases of gene duplications, copies are named and coloured with different shades. For details on number of individuals and exact values, see Supplementary Table. (B) Ontogenetic changes of rod/cone opsin gene expression, and to it related shifts in expression of phototransduction cascade genes GNAT1 (rod-specific) and GNAT2 (cone-specific) for selected teleost taxa. Highlighted in green are special cases of the two aulopiform species that exhibit a discordance between the dominating opsin type (rod-specific) and phototransduction cascade genes (cone-specific) in adults (Lupše et al. 2021).

**Table 1:** Statistical comparison between the younger and older developmental changes for 32 ray-finned fish genera. Summary of beta regression models specifically aimed at proportional data sets (opsin expression as a dependent variable from developmental stage) with the obtained p-values. Alpha levels of significance after the Bonferroni correction additionally marked as equivalent to: < 0.001*** and < 0.05*.

| opsin gene(s) | p-value |
| --- | --- |
| *SWS1* | 0.005 * |
| *SWS2* | 0.040 |
| *RH2* | 0.469 |
| *LWS* | 0.675 |
| rods | 1.6e-09 *** |
| cones | 7.6e-10 *** |
| single cones | 0.950 |
| double cones | 0.302 |

In the selected taxa (Fig. 3), we have specifically focused on the rod vs. cone identity by quantifying the expression of the phototransduction cascade gene *GNAT1* or *GNAT2*, respectively. We found correspondence between the expression of phototransduction cascade gene type and the opsin type (i.e. cone *SWS1*, *SWS2*, *RH2*, *LWS* and rod *RH1*), and detected a clear increase of *GNAT1*:*GNAT2* ratio with ageing, with the exception of the Aulopiformes deep-sea fishes. In this group, a discordance between the dominating opsin type (rod-specific) and phototransduction cascade genes (cone-specific) in adults challenges the rod vs. cone identity and suggests a presence of possibly partially transmuted photoreceptors, potentially similar but not identical to other vertebrates (snakes and geckoes: Simoes et al. 2016; Schott et al. 2019; deep-sea fishes: de Busserolles et al. 2017, Wagner et al. 2019, Lupše et al. 2021; salamanders: Mariani 1986). The overall intriguing visual system of aulopiforms, hence, definitely needs to be investigated further and in more detail (Fig 3, Lupše et al. 2021).

### Developmental switch of the short-wavelength sensitive opsin genes

A trend of age-related shifts in expression also appears within cone opsins (Table 1). Our data set shows a clear decrease in proportional expression of the ultraviolet or UV-sensitive *SWS1* (λ_max_: 347–383 nm) with age (*p* = 0.005; Table 1). Although *SWS1* expression is usually low, it seems to be expressed more in early stages throughout the phylogeny (Fig 1, Table 1). On one hand, UV radiation can result in larval mortality; to mitigate negative effects of exposure, UV avoidance through detection of ultraviolet light and adjustments of vertical position is expected (Ylönen et al. 2004, Guggiana-Nilo and Engert 2016). On the other hand, distinguishing wavelengths belonging to the UV part of the visual spectrum aids younger individuals that feed on zooplankton (Browman et al. 1994, Flamarique et al. 2013, Fattah Ibrahim et al. 2015). With ageing and a shift in diet, UV opsin expression might become irrelevant for some species (Britt et al. 2001), thus potentially explaining why some adults do not express *SWS1* (e.g. *Naso brevirostris*, *Oryzias latipes*), while others still do (e.g. *Danio rerio*, *Poecilia reticulata*, cichlids) (Fig 1, Supp Table). Adult expression of *SWS1*, when seen, seems to play a role in species and/or colour discrimination and mate selection (guppies: Smith et al. 2002; damselfishes: Siebeck et al. 2010; cichlids: Carleton et al. 2016), male aggression (sticklebacks: Rick and Bakker 2008) or is associated with migration events (salmonids: Allison et al., 2006). The blue sensitive *SWS2* cone opsin (λ_max_: 397–482 nm) expression generally increases with age and generally replaces the SWS1 gene in the single cones (Figs 1 and 2, Supp Table Table 1). Interestingly, while some fish (e.g., sturgeons and cyprinids) seem to ontogenetically decrease the proportion of both *SWS1* and *SWS2* opsins, other fish groups (e.g., cichlids) replace one type by another (Fig 3). This switch in single cone opsin expression between *SWS1* and *SWS2* has been shown before e.g., by Spady et al. (2006) in Nile tilapia or by Cheng and Flamarique (2007) in rainbow trout, and it mostly keeps the total single cone opsin expression similar between different developmental stages (Fig. 2).

### Middle and long-wavelength sensitive opsins in double cones

The ontogenetic switch in expression occurs also between the green-sensitive *RH2* (λ_max_: 452–537 nm) and the red-sensitive *LWS* (λ_max_: 501–573 nm) cone opsin types; plus switching between different *RH2* copies also commonly occurs (Fig 3). Values for these typically double-cone opsins vary considerably across the fish phylogeny, albeit a possible weak general trend of a decrease in relative expression of *RH2*, and an increase of *LWS* with age is noticable (Figs 1 and 2, Supp Table; not significant – Table 1), except for groups that completely lost the *LWS* opsin gene. In general, medium-wavelength opsins appear to be of use to all stages (Figs 1 and 2, Supp Table), perhaps due to general presence of corresponding wavelengths in most habitats. Our overview data further seem to show that freshwater species exhibiting the dominance of red-sensitive *LWS* opsin gene expression, whereas in marine species, green-sensitive *RH2* gets to be more dominant (with exceptions) (Fig 1). Namely, for species inhabiting the spectrally narrower deep sea at least during certain parts of their lives (Stomiiformes, Aulopiformes, Trachichthyiformes, Anguilliformes, Gadiformes), *RH2* seems to be the most important, if not the only cone opsin expressed (Fig 1, Lupše et al. 2021). On the other hand, expression of *LWS* in adults might be a response to inhabiting freshwater habitats, such as turbid rivers and murky, eutrophic lakes (e.g., Lake Victoria) where usually, longer wavelengths penetrate to greater depths and are the most prevalent colour of the ambient light (Hofmann et al. 2009, Carleton et al. 2016). Expression of *LWS* is also beneficial for foraging in herbivorous reef fishes, providing them with the visual ability to discriminate benthic algae from coral reef backgrounds (Marshall et al. 2003, Stieb et al. 2017). In some cases, increased *LWS* expression and expanded *LWS* repertoires might also be explained by sexual selection (e.g. in Poeciliidae), where females evolved mate preferences for red and orange male coloration (Watson et al. 2011).

### Age-specific cone opsin gene copies in the selected taxa

We have specifically focused and de-novo sequenced retina transcriptomes of larvae/juveniles and adults of 14 actinopterygian species belonging to five orders spanning the ray-finned fish phylogeny. Apart from the aforementioned rod vs. cone identity assessed by *GNAT* genes, we have additionally focused on switches between copies of the same opsin type in the selected taxa (Fig 3, Supp Table). Namely, we studied the visual opsin gene repertoire in two basal non-teleost fish groups, bichirs (Polypteriformes) and sturgeons (Acipenseriformes), and in teleost riverine cyprinids (Cypriniformes, Ostariophysi), crater-lake cichlids (Cichliformes, Euteleostei) and deep-sea pearleyes and sabretooths (Aulopiformes, Euteleostei). The overall expression patterns are in most cases in accord with the general patterns discussed above (Figure 3, Supp Table), with exceptions seen in the deep-sea fishes (based on our earlier data from Lupše et al. 2021).

In all species but the bichir, we found multiple copies within at least one opsin gene type, namely within the rod *RH1* opsin, and cone *SWS2* and *RH2* opsins. In some species (cyprinids, sturgeon, *Scopelarchus* spp.) we found simultaneous expression of two rod *RH1* copies (Fig 1, Supp Table). All three groups possess the two *RH1* genes in their genome resulting from three independent ancestral gene duplication events (Musilova et al. 2021, Lupše et al. 2021). The *RH1* gene duplicates were lost in the later evolution of the euteleost crown group, and hence most teleost species carry only one *RH1* copy, a phenomenon similar to that seen in “non-fish” vertebrates. These *RH1* copies do not show any sign of ontogenetic switch in studied species, as known e.g. for eels (Hope et al., 1998). On the other hand, we detected several cases of stage-specific copies within cone opsin genes. While *Acipenser ruthenus* and *Abramis brama + Vimba vimba* express only one *SWS2* copy, cichlids express two different *SWS2* genes (Fig 3, Supp Table); this corresponds to multiple copies found in their genome due to the neoteleost- and percomorph-specific *SWS2* gene duplications (Cortesi et al. 2015). Most examined species show an expanded *RH2* repertoire (Fig 3, Supp Table) and the existence of clearly larval and adult-specific copies has been observed in cyprinids, cichlids and in the deep-sea aulopiforms (Fig. 3). Expression of multiple copies might enhance colour vision by increased spectral resolution useful in a particular environment, however reasons for these opsin switches are not yet completely understood. The presence of such stage-specific copies means that species adjust their vision to differing light environments not only through a change in opsin class expression, but also through preferential expression of opsin copies within a single class. In cichlids, a group for which the development of visual system is probably best understood, a shift to longer-wavelength copies is generally observed within a single opsin type (*RH2A* copies replacing *RH2B* with age) or among single-cone opsins (*SWS2* replacing *SWS1*) and has been reported before for different groups of cichlids (e.g., Malawi, Carleton et al. 2008; Nile tilapia, Spady et al., 2006).

Mesopelagic deep-sea aulopiform species have a limited repertoire of cone opsin classes that reflects living in photon-depleted depths (Musilova et al. 2019, Lupše et al. 2021). *Scopelarchus* spp. and *Coccorella atlantica* express only one cone opsin class, namely *RH2* (Fig 3, Supp Table). However, both expanded their *RH2* repertoires and express larval- and adult-specific copies that are thought to be functionally different and most likely best respond to different wavelengths shallow-water epipelagic larvae and mesopelagic deep-water adults encounter (Fig 3, Supp Table) (Lupše et al. 2021). Genomic analyses by Lupše et al. (2021) reveal a total of three, and seven *RH2* cone opsin copies within the genomes of *Coccorella atlantica* and *Scopelarchus michaelsarsi*, respectively. Mesopelagic fish lineages in some cases expand rod opsin repertoires, which are better suited for dim-light conditions (Musilova et al. 2019, Lupše et al. 2021). *Coccorella* and *Scopelarchus*, however, seem to inhabit relatively shallower and photon-richer depths than some other deep-sea fishes, such as Stomiiformes, and might thus benefit also from having extra copies of cone opsins (Lupše et al. 2021).

We have collected a robust data set combining not only our own, but also publicly available genetic data, deposited in databases. This allowed us to detect shared vs. specific expression patterns among different fish groups. We are aware that the collected data set has certain limitations and that many factors could not be controlled in this study. For example, this data set is highly dependent on publicly available material, so there is no control over several potentially relevant factors, such as the sampling conditions, intraspecific variability, other tissues sequenced together with the eyes (as in the entire embryos), etc. Since not all stages are available for all species, we do not present any typical “developmental time series” but rather snapshots of embryos, larvae, juveniles and adults; consequently, more subtle or time-restricted expression patterns could not be detected here. For the purpose of statistical analyses, we have restricted the public data set only to species with two (or more) stages found (Figure 2). To complement the public data we also include our own, controlled data in more detail (Figure 3). Despite certain limitations, our combined dataset provides robust evidence for expression patterns shared across distantly related fish groups, as it highlights general trends, and more detailed conclusions achieved through in-detail analyses of species specifically sequenced within this study.

## Conclusions

To conclude, this study aims to identify general patterns of the visual opsin gene expression shared among ray-finned fishes, and to detect similarities in the ontogenetic changes between opsin gene types. We found that the rod:cone opsins ratio increased with age in fish species, supporting the conserved cone-to-rod developmental pathway. We also noted the increased importance of the long-wavelength sensitive *LWS* opsin genes, and the decreased importance of the short-wavelength sensitive *SWS1* opsin gene, observed across ray-finned fish phylogeny (e.g. in sturgeons, cyprinids and cichlids). We have further detected the existence of different stage-specific *RH2* copies, which are switched during development. To conclude, fish visual systems are evolutionary and developmentally very dynamic and future studies focused on particular fish groups promise to throw further light on exact mechanisms, patterns and reasons for this extreme sensory system diversity.

## Acknowledgements

We would like to express our deep gratitude to Adrian Indermaur and Petra Horká for their help with the field sampling. We thank local people of the Barombi and Bemé village to allow us to fish in the Barombi Mbo and Bermin lakes, and the Ministry of Scientific Research and Innovation in Cameroon to grant us research permits. NL and ZM were supported by the Swiss National Science Foundation (PROMYS – 166550), ZM by the PRIMUS Research Programme (Charles University), the Czech Science Foundation (21-31712S) and the Basler Stiftung fuer Experimentelle Zoologie. VT and PK were supported by GAUK (1767618). VK was funded by Ministry of Education, Youth and Sports of the Czech Republic – the project Reproductive and Genetic Procedures for Preserving Fish Biodiversity and Aquaculture (CZ.02.1.01/0.0/0.0/16_025/0007370).

## References

Allison WT, Dann SG, Veldhoen KM, Hawryshyn CW. 2006 Degeneration and regeneration of ultraviolet cone photoreceptors during development in rainbow trout. Journal of Comparative Neurology, 499(5), 702–715.

Andrews S. 2017. FastQC: a quality control tool for high throughput sequence data. 2010.

Betancur-R R. et al. 2017 Phylogenetic classification of bony fishes. BMC evolutionary biology, 17(1), 162

Blaxter JHS. 1986 Development of sense organs and behaviour of teleost larvae with special reference to feeding and predator avoidance. Transactions of the American Fisheries Society, 115,98–114

Browman HI, Flamarique IN, Hawryshyn CW. 1994 Ultraviolet photoreception contributes to prey search behavior in two species of zooplanktivorous fishes. J. Exp. Biol. 186: 187–198.

Carleton KL, Dalton BE, Escobar-Camacho D, Nandamuri SP. 2016 Proximate and ultimate causes of variable visual sensitivities: insights from cichlid fish radiations. Genesis, 54(6), 299–325.

Carleton KL, Escobar-Camacho D, Stieb SM, Cortesi F, Marshall NJ. 2020. Seeing the rainbow: mechanisms underlying spectral sensitivity in teleost fishes. Journal of Experimental Biology, 223(8).

Carleton KL, Kocher TD. 2001 Cone opsin genes of African cichlid fishes: tuning spectral sensitivity by differential gene expression. Molecular Biology and Evolution, 18(8), 1540–1550.

Carleton KL, Spady TC, Streelman JT, Kidd MR, McFarland WN, Loew ER. 2008 Visual sensitivities tuned by heterochronic shifts in opsin gene expression. BMC Biology, 6(1), 1–14.

Carvalho PS, Noltie DB, Tillitt DE. 2002 Ontogenetic improvement of visual function in the medaka *Oryzias latipes* based on an optomotor testing system for larval and adult fish. Animal Behaviour, 64(1), 1–10.

Chang CH, Catchen J, Moran RL, Rivera-Colón AG, Wang YC, Fuller RC. 2021 Sequence analysis and ontogenetic expression patterns of cone opsin genes in the bluefin killifish (*Lucania goodei*). Journal of Heredity. 112(4), 357–366.

Chang CH, Wang YC, Shao YT, Liu SH. 2020 Phylogenetic analysis and ontogenetic changes in the cone opsins of the western mosquitofish (*Gambusia affinis*). PloS one, 15(10), e0240313.

Cheng CL, Flamarique IN. 2007 Chromatic organization of cone photoreceptors in the retina of rainbow trout: single cones irreversibly switch from UV (*SWS1*) to blue (*SWS2*) light sensitive opsin during natural development. Journal of Experimental Biology. 210(23), 4123–4135

Cortesi F, et al. 2021. Multiple ancestral duplications of the red-sensitive opsin gene (LWS) in teleost fishes and convergent spectral shifts to green vision in gobies. bioRxiv: https://doi.org/10.1101/2021.05.08.443214.

Cortesi F et al. 2016 From crypsis to mimicry: changes in colour and the configuration of the visual system during ontogenetic habitat transitions in a coral reef fish. Journal of Experimental Biology, 219(16), 2545–2558.

Cortesi F, et al. 2015 Ancestral duplications and highly dynamic opsin gene evolution in percomorph fishes. Proceedings of the National Academy of Sciences, 112(5), 1493–1498.

Cottrill PB, Davies WL, Bowmaker JK, Hunt DM, Jeffery G. 2009 Developmental dynamics of cone photoreceptors in the eel. BMC Developmental Biology, 9(1), 1–9.

Cribari-Neto F & Zeileis A. 2010. Beta regression in R. Journal of statistical software, 34, 1–24.

De Busserolles F, Marshall NJ, Collin SP. 2014. The eyes of lanternfishes (Myctophidae, teleostei): novel ocular specializations for vision in dim light. Journal of Comparative Neurology, 522(7), 1618–1640.

de Busserolles F, et al. 2017 Pushing the limits of photoreception in twilight conditions: The rod-like cone retina of the deep-sea pearlsides. Science advances, 3(11), eaao4709.

Evans BI, Browman HI 2004 Variation in the development of the fish retina. In American Fisheries Society Symposium (Vol. 40, pp. 145–166).

Evans BI, Fernald RD. 1990 Metamorphosis and fish vision. Journal of neurobiology, 21(7), 1037–1052.

Fattah Ibrahim A, Castilho Noll MSM, Valenti WC. 2015. Zooplankton capturing by Nile Tilapia, *Oreochromis niloticus* (Teleostei: Cichlidae) throughout post-larval development. Zoologia (Curitiba) 32:469–475.

Fernald RD. 1985 Growth of the teleost eye: novel solutions to complex constraints. Environmental Biology of Fishes, 13(2), 113–123.

Flamarique IN. 2013 Opsin switch reveals function of the ultraviolet cone in fish foraging. Proceedings of the Royal Society B: Biological Sciences, 280(1752), 20122490.

Fogg LG, Cortesi F, Lecchini D, Gache C, Marshall NJ, De Busserolles F. 2022. Development of dim-light vision in the nocturnal reef fish family Holocentridae I: retinal gene expression. Journal of Experimental Biology.

Fuiman LA. 1993 Development of predator evasion in Atlantic herring, *Clupea harengus* L. Animal Behaviour, 45(6), 1101–1116.

Fuiman LA, Delbos BC. 1998 Developmental changes in visual sensitivity of red drum, *Sciaenops ocellatus*. Copeia, 936–943.

Guggiana-Nilo DA, Engert F. 2016 Properties of the visible light phototaxis and UV avoidance behaviors in the larval zebrafish. Frontiers in behavioral neuroscience, 10, 160.

Hagedorn M, Fernald RD. 1992. Retinal growth and cell addition during embryogenesis in the teleost, *Haplochromis burtoni*. Journal of Comparative Neurology, 321(2), 193–208.

Härer A, Torres-Dowdall J, Meyer A. 2017. Rapid adaptation to a novel light environment: the importance of ontogeny and phenotypic plasticity in shaping the visual system of Nicaraguan Midas cichlid fish (*Amphilophus citrinellus* spp.). Molecular ecology, 26(20), 5582–5593.

Helfman G, Collette BB, Facey DE, Bowen BW. 2009. The diversity of fishes: biology, evolution, and ecology (John Wiley & Sons: West Sussex, UK).

Helvik JV, Drivenes Ø, Harboe T, Seo HC. 2001 Topography of different photoreceptor cell types in the larval retina of Atlantic halibut (*Hippoglossus hippoglossus*). Journal of Experimental Biology, 204(14), 2553–2559.

Hofmann CM, O’Quin KE, Marshall NJ, Cronin TW, Seehausen O, Carleton KL. 2009. The eyes have it: regulatory and structural changes both underlie cichlid visual pigment diversity. PLoS biology, 7(12), e1000266.

Hope A, Partridge J, Hayes P. 1998. Rod opsin shifts in the European eel, Anguilla anguilla (L). Proc. R. Soc. Lond B, 265(January), 869–874.

Hunt DM, Hankins MW, Collin SP, Marshall NJ. 2014. Evolution of visual and non-visual pigments (Vol. 4). Boston, MA: Springer.

Hunt DM, Fitzgibbon J, Slobodyanyuk SJ, Bowmaker JK, Dulai KS. 1997 Molecular evolution of the cottoid fish endemic to Lake Baikal deduced from nuclear DNA evidence. Molecular Phylogenetics and Evolution, 8(3), 415–422.

Hunter JR, Coyne KM. 1982 The onset of schooling in northern anchovy larvae, *Engraulis mordax*. CalCOFI Rep, 23, 246–251.

Kearse M, et al. 2012. Geneious Basic: an integrated and extendable desktop software platform for the organization and analysis of sequence data. Bioinformatics, 28(12), 1647–1649.

King JR, McFarlane GA. 2003. Marine fish life history strategies: applications to fishery management’, Fisheries Management and Ecology, 10, 249–64

La Vail MM, Rapaport DH, Rakic P. 1991. Cytogenesis in the monkey retina. J. Comp. Neurol. 309, 86–114. doi:10.1002/cne.903090107.

Lamb TD. 2013. Evolution of phototransduction, vertebrate photoreceptors and retina. Progress in retinal and eye research, 36, 52–119.

Lupše N, et al. 2021 Visual gene expression reveals a cone to rod developmental progression in deep-sea fishes. Molecular Biology and Evolution 38(12):5664–5677

Magnuson JT, Stieglitz JD, Garza SA, Benetti DD, Grosell M, Roberts AP. 2020 Development of visual function in early life stage mahi-mahi (*Coryphaena hippurus*). Marine and Freshwater Behaviour and Physiology, 53(5-6), 203–214.

Malinsky M, et al. 2015 Genomic islands of speciation separate cichlid ecomorphs in an East African crater lake. Science. 350(6267), 1493–8. http://doi.org/10.1126/science.aac9927

Manousaki et al. 2013 Parsing parallel evolution: ecological divergence and differential gene expression in the adaptive radiations of thick-lipped Midas cichlid fishes from Nicaragua. Molecular ecology, 22(3), 650–669.

Mariani AP, Boycott BB. 1986. Photoreceptors of the larval tiger salamander retina. Proceedings of the Royal Society of London. Series B. Biological Sciences, 227: 483–92.

Marshall NJ, Jennings K, McFarland WN, Loew ER, Losey GS. 2003 Visual biology of Hawaiian coral reef fishes. III. Environmental light and an integrated approach to the ecology of reef fish vision. Copeia, 2003(3), 467–480.

Mears AJ, et al. 2001. *Nrl* is required for rod photoreceptor development. Nature genetics, 29(4), 447–452.

Musilova Z, et al. 2019a. Vision using multiple distinct rod opsins in deep-sea fishes. Science, 364(6440), 588–592.

Musilova Z, Salzburger W, Cortesi F. 2021. The visual opsin gene repertoires of teleost fishes: evolution, ecology, and function. Annual review of cell and developmental biology, 37, 441–468.

Musilova Z, Cortesi F. 2021 Multiple ancestral and a plethora of recent gene duplications during the evolution of the green sensitive opsin genes (*RH2*) in teleost fishes. bioRxiv.

Musilova, et al. 2019 Evolution of the visual sensory system in cichlid fishes from crater lake Barombi Mbo in Cameroon. Molecular Ecology 5010–5031. http://doi.org/10.1111/mec.15217

Pankhurst PM, Pankhurst NW, Montgomery JC. 1993. Comparison of behavioural and morphological measures of visual acuity during ontogeny in a teleost fish, *Forsterygion varium*, Tripterygiidae (Forster, 1801). Brain, Behavior and Evolution, 42(3), 178–188.

Pointer MA, Carvalho LS, Cowing JA, Bowmaker JK, Hunt DM. 2007. The visual pigments of a deep-sea teleost, the pearl eye Scopelarchus analis. Journal of Experimental Biology, 210(16), 2829–2835.

Raymond PA. 1995. Development and morphological organization of photoreceptors. Neurobiology and Clinical Aspects of the Outer Retina (pp. 1–23). Springer, Dordrecht.

Ricci V, Ronco F, Musilova Z, Salzburger W. 2022. Molecular evolution and depth-related adaptations of rhodopsin in the adaptive radiation of cichlid fishes in Lake Tanganyika. Molecular Ecology

Rick IP, Bakker TC. 2008 Males do not see only red: UV wavelengths and male territorial aggression in the three-spined stickleback (*Gasterosteus aculeatus*). Naturwissenschaften, 95(7), 631–638.

Roux N et al. 2022. The multi-level regulation of clownfish metamorphosis by thyroid hormones, bioRxiv, 10.1101/2022.03.04.482938.

Savelli I, Novales Flamarique I, Iwanicki T, Taylor JS. 2018. Parallel opsin switches in multiple cone types of the starry flounder retina: tuning visual pigment composition for a demersal life style. Scientific Reports, 8, 4763.

Schott RK, Bhattacharyya N, Chang BS. 2019. Evolutionary signatures of photoreceptor transmutation in geckos reveal potential adaptation and convergence with snakes. Evolution, 73(9), 1958–1971.

Sernagor E, Eglen S, Harris B, Wong R. 2006. Retinal development. Cambridge University Press.

Shand J, Hart NS, Thomas N, Partridge JC. 2002. Developmental changes in the cone visual pigments of black bream *Acanthopagrus butcheri*. Journal of Experimental Biology, 205(23), 3661–3667.

Shen YC, Raymond PA. 2004. Zebrafish cone-rod (*crx*) homeobox gene promotes retinogenesis. Developmental biology, 269(1), 237–251.

Siebeck UE, Parker AN, Sprenger D, Mäthger LM, Wallis G. 2010 A species of reef fish that uses ultraviolet patterns for covert face recognition. Current Biology, 20(5), 407–410.

Simoes BF et al. (2016). Multiple rod–cone and cone–rod photoreceptor transmutations in snakes: evidence from visual opsin gene expression. Proceedings of the Royal Society B: Biological Sciences, 283(1823), 20152624.

Smith EJ, Partridge JC, Parsons KN, White EM, Cuthill IC, Bennett AT, Church SC. 2002 Ultraviolet vision and mate choice in the guppy (*Poecilia reticulata*). Behavioral Ecology, 13(1), 11–19.

Stieb SM, Cortesi F, Sueess L, Carleton KL, Salzburger W, Marshall NJ. 2017 Why UV vision and red vision are important for damselfish (Pomacentridae): structural and expression variation in opsin genes. Molecular ecology, 26(5), 1323–1342.

Stieb SM, Carleton KL, Cortesi F, Marshall NJ, Salzburger W. 2016 Depth-dependent plasticity in opsin gene expression varies between damselfish (Pomacentridae) species. Molecular ecology, 25(15), 3645–3661.

Sugawara T, et al. 2005 Parallelism of amino acid changes at the *RH1* affecting spectral sensitivity among deep-water cichlids from Lakes Tanganyika and Malawi. Proceedings of the National Academy of Sciences, 102(15), 5448–5453.

Tettamanti V, de Busserolles F, Lecchini D, Marshall NJ, Cortesi F. 2019 Visual system development of the spotted unicornfish, *Naso brevirostris* (Acanthuridae). Journal of Experimental Biology, 222(24).

Valen R, et al. 2016 The two-step development of a duplex retina involves distinct events of cone and rod neurogenesis and differentiation. Developmental biology, 416(2), 389–401.

Valen R, Karlsen R, Helvik JV. 2018 Environmental, population and life-stage plasticity in the visual system of Atlantic cod. Journal of Experimental Biology, 221(1), jeb165191.

Wagner HJ, Partridge JC, Douglas RH. 2019. Observations on the retina and ‘optical fold’of a mesopelagic sabretooth fish, *Evermanella balbo*. Cell and tissue research, 378(3), 411–425.

Watson CT, et al. 2011. Gene duplication and divergence of long wavelength-sensitive opsin genes in the Guppy, *Poecilia reticulata*. J Mol Evol. 72(2):240–252.

Ylönen O, Heikkilä J, Karjalainen J. 2004 Metabolic depression in UV-B exposed larval coregonids. Annales Zoologici Fennici (pp. 577–585).

Yokoyama S. 2008 Evolution of dim-light and color vision pigments. Annu. Rev. Genomics Hum. Genet., 9, 259–282.

Yokoyama S, Jia H. 2020 Origin and adaptation of green-sensitive (*RH2*) pigments in vertebrates. FEBS Open Bio, 10(5), 873–882.

